# Resistance to different antibiotics in clinical isolates of *Enterococcus faecalis* obtained from urine cultures of patients with urinary tract infection

**DOI:** 10.1101/2020.09.20.305623

**Authors:** Mariana Islas Rodríguez, José Carlos Valencia Esquivel, Silvia Patricia Rodríguez Peña, Elisangela Oliveira de Freitas, Jorge Angel Almeida Villegas

## Abstract

**Objective:** To identify patterns of resistance against various antibiotics in *Enterococcus faecalis* in urinary tract infections in a population of the Toluca valley, Mexico

**Methods:** 155 samples were collected from patients with suspected urinary tract infection without exclusion criteria such as age or gender. Automated equipment was used for the identification of the etiological agent and sensitivity tests.

**Results:** 80 positive cultures were obtained, of which 20 strains belong to Enterococcus faecalis, which show 100% sensitivity for penicillins, linezolid, vancomycin, bacitracin, a high pattern of sensitivity for quinolones, and a high pattern of resistance to rifampicin, erythromycin and 100% resistance in tetracycline

**Conclusion:** It shows 100% sensitivity to penicillins, vancomycin and linezolid, first-line treatments and for cases of infection complicated by Enterococci. And 100% resistance for tetracycline and high resistance patterns for erythromycin and rifampin.

## Introduction

Urinary tract infections (UTIs) are the second leading cause of community infection causing hospital admission.^1^ Urinary tract infections are especially common among women, the vast majority of whom experience at least one episode of infection in their lifetime. A significant subset (25–40%) of women also have recurrent urinary tract infections (UTIs), with multiple infections that recur over months or years, in some cases. Other clinical problems related to urinary infection include asymptomatic bacteriuria (AB) and patients with complicated urinary tract infection (urinary infection). Nosocomial urinary tract infection (generally a reflection of catheter-associated infections) constitutes about 20-30% of all hospital-acquired infections and are common sources of nosocomial bacteremia.^2^ The diagnosis of a urinary infection can be made based on a combination of symptoms and a positive urine culture or analysis.^3^

Enterococcal species are core constituents of the intestinal flora of many animal species ranging from humans to flies. Enterococci have gained notoriety over the past few decades as frequent causesof multiple antibiotic resistant, hospital-acquired bloodstream,urinary tract and surgical wound infections; and because of theircapacity to transfer antibiotic resistances to other microbes.^4^ Grampositive bacterium *Enterococcus faecalis,which* occupy overlapping niches in the normal mammalian microbiome, including in the gastrointestinal tract, oral cavity,and urogenital tract.^5^

The incidence of UTIs due to *Enterococcus faecalis* has risen steadily over the years, and infections due to multiple-drug-resistant strainspresent a significant medical problem and are a common cause of chronic or recurrent UTIs, especially those associated with structural abnormalities and instrumentation.^6^

*Enterococcus faecalis* UTIsc are of particular concernin that they are intrinsically resistant to first-line antimicrobial agents, especially vancomycin.^7^

Due to the rapid adaptation of bacteria, antimicrobial treatment for the management of urinary tract infections is becoming increasingly difficult, causing the antimicrobial profile of bacterial uropathogens to change over the years, despite having better antimicrobial agents. Resistance to these has increased significantly due to their incorrect use, limiting therapeutic options.^8^ Currently several bacterial strains are resistant to practically all known antibiotics. Such as carbapenems, cephalosporins, macrolides, and penicillins.^9^

## Methods

In the present study, 155 urine cultures were performed in patients with clinical suspicion of urinary tract infection, without exclusion criteria such as gender or age. In a town in the center of the Toluca Valley, Mexico.

Of the 155 urine cultures performed, the distribution was; 107 people of the female gender representing 69% of the study population and 48 men representing 31%. The age ranges in the lower limit is a 1-year-old female patient and for the higher limit is a 91-year-old patient, with an average of 30 years.

For the study, 100 ml of the first urine of the day were collected with the necessary hygiene conditions for the collection of the specimen.

An aliquot was taken with a calibrated handle, and blood agar was inoculated to perform the colony count and chromogenic agar. Both Petri dishes were incubated for 24 hours at 37°C.

A colony isolated from the chromogenic agar medium was taken and resuspended in diluent medium, which was placed on the plates of the automated equipment.

The identification of the etiological agent as well as the sensitivity tests were performed using an automated method using the walkaway SI 96 from beckman coulter equipment.

## Results and Discussion

Of the urine cultures obtained and performed, 80 of them were positive for the development of microorganisms, which represents 51.6%, on the other hand, 48.4% representing 75 cultures that did not present microbial development.

20 *Enterococcus faecalis* strains were obtained, representing 25% of the total positive cultures for microbial development. For the evaluated antibiotics, marked resistance patterns were obtained for certain relatives, such as 100% sensitivity in beta-lactams (penicillin and ampicillin), and other antibiotics whose mechanism of action is directed to the synthesis of the cell wall such as vancomycin, daptomycin or linezolid. For the quinolones levofloxacin and ciprofloxacin, there is a sensitivity of 90% for each respectively. In these antibiotics little resistance is shown, however in the case of nitrofurans there are 25% resistant strains, 25% strains with intermediate sensitivity and only the remaining 50% are sensitive. Rifampicin has 50% resistant strains and 15% strains with intermediate sensitivity, for the macrolide erythromycin a resistance of 75% is observed and finally 100% resistance for tetracycline. Table 1

**Table 1.** Antibiotic resistance patterns in Enterococcus faecalis.

|  | Resistance patterns |  |  |
| --- | --- | --- | --- |
|  | Resistant | Intermediate sensitivity | Sensitive |
| Ampicillin | 0 | 0 | 20 |
| Ciprofloxacin | 2 | 0 | 18 |
| Daptomycin | 0 | 0 | 20 |
| Erythromycin | 15 | 0 | 5 |
| Levofloxacin | 1 | 1 | 18 |
| Linezolid | 0 | 0 | 20 |
| Penicillin | 0 | 0 | 20 |
| Rifampicin | 10 | 3 | 7 |
| Tetracycline | 20 | 0 | 0 |
| Vancomycin | 0 | 0 | 20 |
| Nitrofurans | 5 | 5 | 10 |

Graph number 1 shows the resistance patterns of the 20 isolates of enterococcus facecalis and their high resistance index in tetracycline, erythromycin and rifampicin, and their high sensitivity to pinicillins, daptomycin, linezolid and vancomycin.

**Graph 1.**
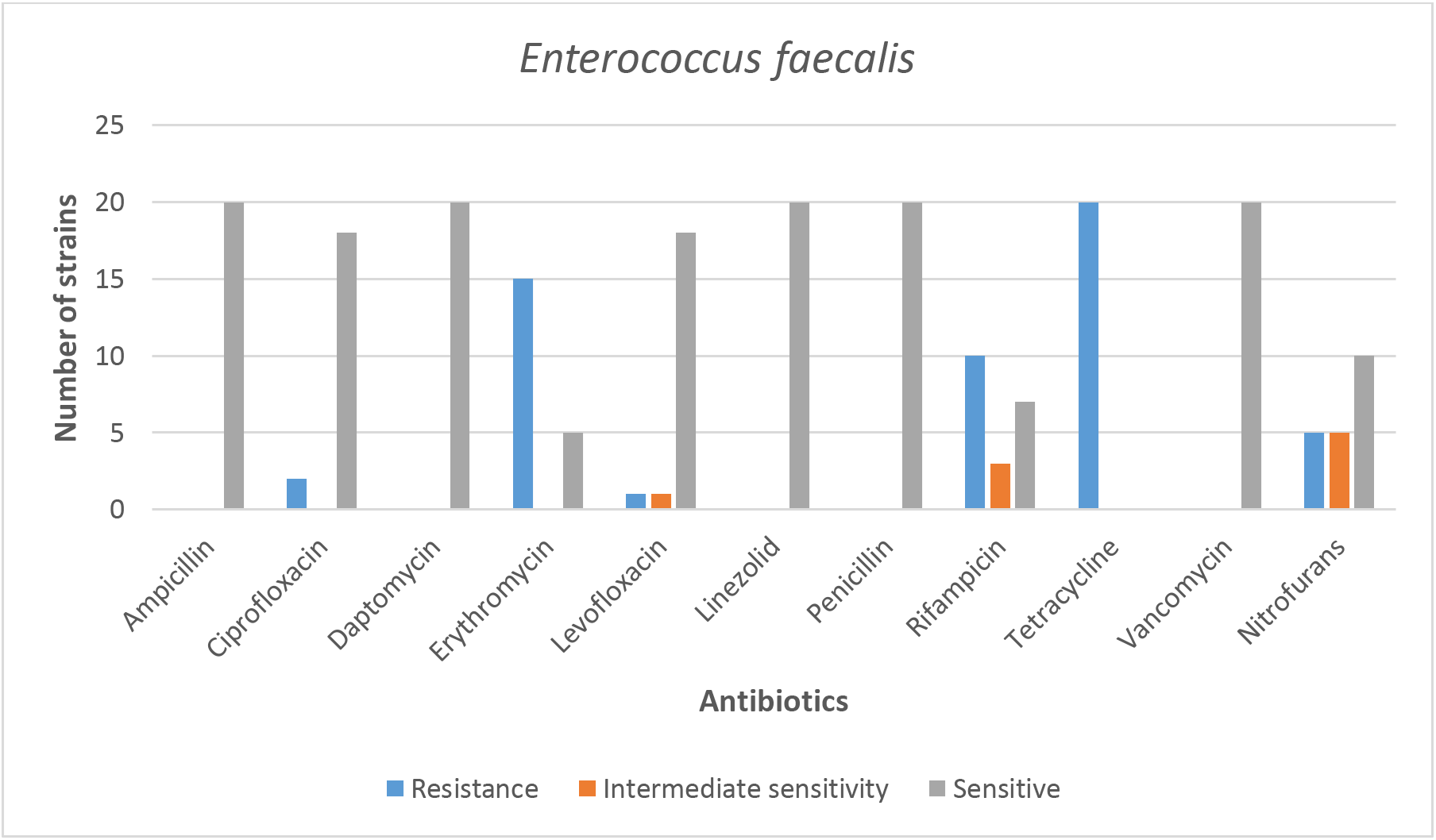
Antibiotic resistance patterns in *Enterococcus faecalis*.

There are three main mechanisms of resistance to the family of macrolides, which make up a triad with other macrolide drugs, lincosamides and streptogramins, which are the; modification of the target drug, inactivation of the drug and active exit of the antibiotic. The high number of strains resistant to erythromycin may suggest some of the aforementioned mechanisms, however, the literature indicates the presence of efflux pumps specific for erythromycin, and suggests the greater resistance mechanism for the enterococcal genus, another mechanism could be mediated by erm methylases of the ermBermAM hybridization class that has been described in *Enterococcus* isolates.^12^

In *Enterococci,* two major groups of Tetracycline resistance (tet) genes have been recognized. The first group confers resistance by ribosomal protection (RP) and includes the genes tet(M), tet(O), and tet(S), which have been detected in Enterococcus spp. A second group mediates energy-dependent efflux of TC from cells and is represented *in Enterococci* by the tet(K) and tet(L) genes.^11^

Mechanisms that may explain the high resistance to tetracycline in the clinical isolates of the present study.

## Conclusions

In the present study it was possible to obtain the resistance patterns of enterococcus faecalis, obtained from urine cultures in patients with urinary tract infection in patients from the Toluca Valley, Mexico. It shows 100% sensitivity to penicillins, vancomycin and linezolid, first-line treatments and for cases of infection complicated by Enterococci. And 100% resistance for tetracycline and high resistance patterns for erythromycin and rifampin.

## Ethical approval

The following study was approved by the ethical committee of the Pasteur laboratorios

No permission of the patients was required for the obtention of the clinical isolates since they are no human material and are routinely recovered from patients cultures.

## Financing

No funding was received to carry out this work.

## Acknowledgment

We thank all the institutions that allowed us to carry out this research work.

